# The Microspherule protein 1(MCRS1) homolog interacts with the Myb-like transcription factor DRMY1 and is essential for embryogenesis in *Arabidopsis thaliana*

**DOI:** 10.1101/2023.07.11.548616

**Authors:** Huan Howard Huo, Ming Luo, Yuh-Ru Julie Lee, Bo Liu

**Affiliations:** Department of Plant Biology, University of California, Davis, CA 95616, USA; Biotechnology Research Center, Southwest University, Beibei, Chongqing, China

**Keywords:** MCRS1 (Microspherule protein 1), DRMY1 (Developmentally Regulated Myb-like1), Embryogenesis, Transcription, Arabidopsis

## Abstract

The evolutionarily conserved Microspherule protein 1 (MCRS1) has diverse functions from transcriptional regulation to stabilization of microtubule minus ends in acentrosomal spindles in mammals. A previous study suggested that in the model plant *Arabidopsis thaliana*, inactivation of an MCRS1 homolog gene led to aborted embryogenesis. To test whether this lethality was caused by defects associated with transcription or mitosis, we used the heterozygous *mcrs1* mutant to examine whether the gene was required for mitosis during gametogenesis. Results of reciprocal crosses between the *mcrs1* mutant and the wild-type plant showed that the *MCRS1* gene was dispensable for mitotic cell divisions associated with the development of both male and female gametophytes. An MCRS1-GFP fusion protein was expressed in the *mcrs1* mutant and suppressed the mutation as reported by restored growth. This functional fusion protein exclusively localized to interphase nuclei and became undetectable during mitosis before returning to the reforming daughter nuclei. Affinity purification of the MCRS1-GFP protein resulted in consistent recovery of the Myb-like transcription factor DRMY1 (Developmentally Regulated Myb-like1) but not microtubule-associated factors. The association was further supported by the evidence of a direct interaction in living cells. Hence, the plant MCRS1 was concluded to play a role in the gene transcription in sporophyte development.

## Introduction

The Microspherule protein 1 (MCRS1) is an evolutionarily conserved protein and functions as a critical regulator of cell cycle progression in mammals (Huang et al. 2022). The protein is featured with the characteristic N-terminal MCRS_N domain and a C-terminal forkhead-associated (FHA) domain. In human cells, it interacts with functionally diversified proteins and is often associated with transcriptional regulation of genes involved in cell proliferation by being a component of the INO80 complex involved in ATP-dependent chromatin-remodeling (Jin et al. 2005). Besides, MCRS1 also plays a role in nucleolus sequestering of ribosome-related proteins to promote rRNA transcription and other protein(s) in order to relieve transcriptional repression (Lin and Shih 2002; Shimono et al. 2005).

Surprisingly, other studies also showed that MCRS1 localizes to the spindle poles during early stages of mitosis and plays a critical role in the assembly of acentrosomal spindle (Meunier and Vernos 2011). The protein is recruited to the minus ends of kinetochore fibers, but not centrosomal microtubules, by the epigenetic regulator of KAT8-associated nonspecific lethal (KANSL) complex via direct interaction (Meunier et al. 2015). It stabilizes microtubule minus ends possibly through the regulation of the microtubule depolymerase of kinesin-13(Meunier and Vernos 2011).

In the model plant *Arabidopsis thaliana*, the AT3G54350 locus encodes a protein sharing high degree of homology to the human MCRS1. It was shown recently that inactivation of this *MCRS1* homologous gene caused severe defects in embryogenesis as shown in the *emb1967-1* and *emb1967-2* mutants (Meinke 2020). One might argue that such a lethality in *A. thaliana* was caused by defects in the assembly of acentrosomal spindles during mitosis and/or meiosis if the plant MCRS1 homolog shared the microtubule-associated function as its human counterpart. Alternatively, *MCRS1* may be an essential gene because of its role in transcriptional regulation in the plant. To characterize the specific function of MCRS1, we performed protein localization and purification experiments using plants expressing a functional MCRS1-GFP (green fluorescent protein) fusion protein. We learned from the experimental outcome that MCRS1 was a nuclear factor that associated with the Myb family transcription factor DRMY1 and it did not gain a microtubule-associated function in *A. thaliana*.

## Results

### Identification of MCRS1 homologs in Arabidopsis and other land plants

When the human MCRS1 was compared to proteins in *A. thaliana* (https://www.arabidopsis.org/), the protein encoded by the locus AT3G54350 was found to share a high degree of homology in both the N-terminal MCRS_N domain as well as the C-terminal FHA domain (Supple. Fig. 1 and Fig. 1). Therefore, we named this gene *AtMCRS1* because of such high degrees of homology between AtMCRS1 and HsMCRS1 in both characteristic domains: 59.62% and 37.04% identical in MCRS_N domain and FHA domain, respectively (Fig. 1). To learn whether *AtMCRS1* is conserved in land plants, we searched proteomes of representative land plants, namely the bryophytes of liverwort (*Marchantia polymorpha*) and moss (*Physcomitriella patens*), the early vascular plant lycophyte (*Selaginella moellendorffii*), and the monocot rice (*Oryza sativa*). All these plants produce the MCRS1 homologous proteins with highly conserved N-terminal MCRS_N domain and the C-terminal FHA domain (Supple. Fig. 1). Therefore, studies of the AtMCRS1 gene likely lead to the understanding of common functions of the plant homologs of this conserved protein.

**Figure 1.**
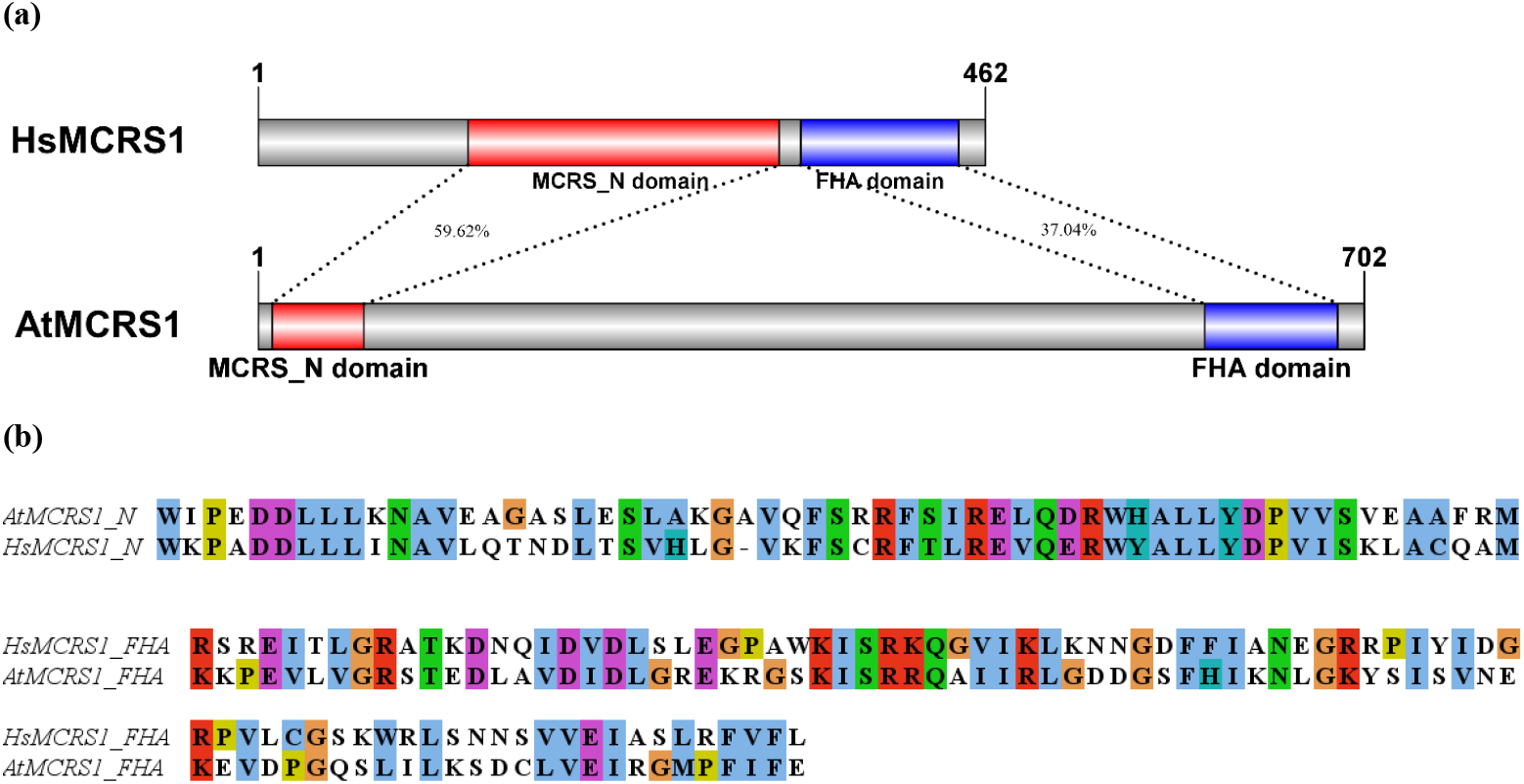
Homology between HsMCRS1 and AtMCRS1. (a) Domain architecture of HsMCRS1 and AtMCRS1. The predicted MCRS_N domains are labeled in red and FHA domains are labeled in blue. The MCRS_N domains share 59.62% sequence identity, and the FHA domains share 37.04% sequence identity. Hs: *Homo sapiens*; At: *Arabidopsis thaliana*; (b) Sequence alignments of the homologous regions of MCRS_N domains and FHA domains. The conserved amino acids are labeled in colours.

### *AtMCRS1* is essential for sporophyte but not gametophyte development

The At3G54350 locus has been annotated as one of the *EMBRYO-DEFECTIVE* (*EMB*) genes and two corresponding T-DNA mutant *emb1967-1* and *emb1967-2* were identified (Meinke et al. 2008; Meinke 2020). Homozygous *emb1967-1* and *emb1967-2* mutants showed early termination of the embryo development before embryos transition to heart shape stage (https://seedgenes.org/). The *emb1967-1* mutation had a T-DNA fragment harboring a glufosinate (Basta) resistant cassette, whereas *emb1967-2* had a T-DNA linked to the kanamycin-resistant gene (Supple. Fig 2)(Meinke et al. 2008; Meinke 2020). Similar to what has been reported, we did not recover any homozygous mutant plants from either heterozygous parent. Aborted seeds were detected in both pools of F2 populations (Fig. 2).

**Figure 2.**
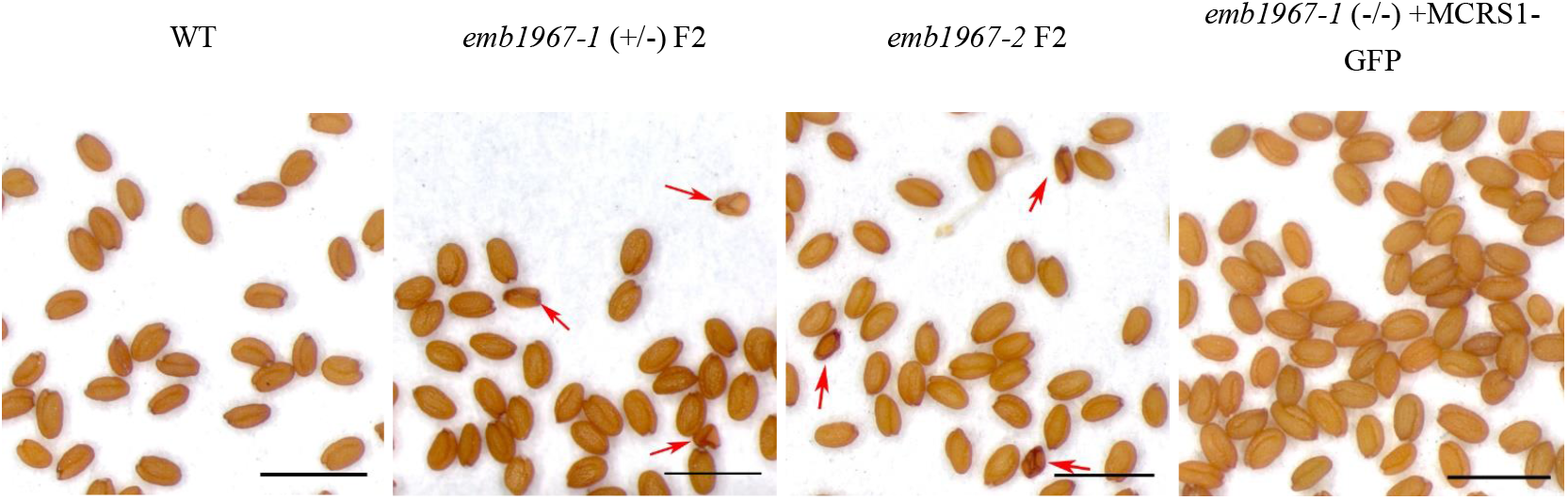
AtMCRS1 is essential for seed development. Complementation of the *emb1967-1* mutation by AtMCRS1-GFP expression under its native promotor as reflected by the suppression of the seed abortion phenotype. Seeds of wild type plants, *mcrs1* mutant plants (Progenies of heterozygous *emb1967-1* and *emb1967-2* parents), and rescued plants (*emb1967-1* (-/-) +AtMCRS1-GFP). Red arrows in the middle two panels indicate aborted and shrunken seeds. Scale bars=2 mm.

The lack of homozygous offspring prompted us to investigate whether the segregation of this mutation during sexual reproduction was defected. Among the plants produced by the heterozygous *emb1967-1* mutant, we found a Basta sensitive-to-resistant ratio of 0.49:1 (Table 1) which matched the 1:2 ratio predicted by the Mendelian Segregation Law. Therefore, the segregation of the mutant allele was not affected.

**Table 1.**
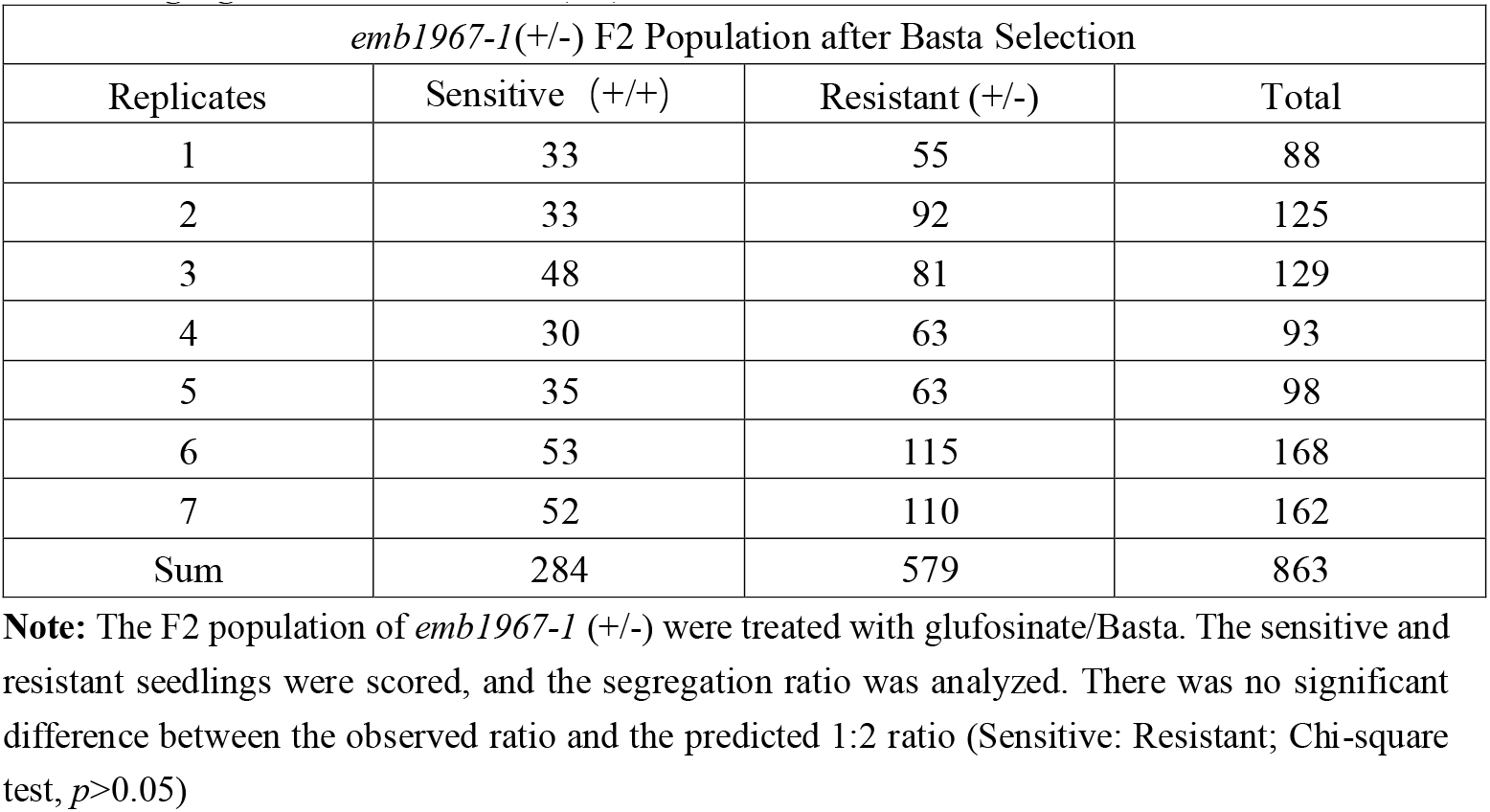
Segregation of *emb1967-1* (+/-) F2 population.

To further test whether the mutations affected gametogenesis that were dependent on mitotic divisions after meiosis, we performed reciprocal crosses between the heterozygous *emb1967-1* mutant and the wild type (Col-0) and examined the segregation ratio of the progenies with the BASTA sensitivity test (Table 2). Ideally, if all gametophytes were functional then a 1:1 segregation ratio would be expected. When the wild type pollen grains were used to pollinate the mutant, a Basta sensitive-to-resistant ratio of 1.13:1 was detected. Conversely, a similar ratio of 1.11:1 was observed when the mutant pollen grains were used to pollinate the wild type stigmas. Therefore, we concluded AtMCRS1 was not required for either male or female gametogenesis and associated mitotic divisions.

**Table 2.** Reciprocal Crosses between the WT plants and *emb1967-1* (+/-)

| WT♂×*emb1967-1* (+/-) ♀ Progenies after Basta Selection |  |  |  |
| --- | --- | --- | --- |
| Replicates | Sensitive (+/+) | Resistant (+/-) | Total |
| 1 | 23 | 29 | 52 |
| 2 | 54 | 39 | 93 |
| Sum | 77 | 68 | 145 |
| WT♀×*emb1967-1* (+/-) ♂ Progenies after Basta Selection |  |  |  |
| Replicates | Sensitive (+/+) | Resistant (+/-) | Total |
| 1 | 11 | 12 | 23 |
| 2 | 60 | 52 | 112 |
| Sum | 71 | 64 | 135 |

**Note:** The progenies from reciprocal crosses between the WT and *emb1967-1* (+/-) were treated with glufosinate. The sensitive and resistant progenies were scored, and the segregation ratio was analyzed. There was no significant difference between the observed ratio and the predicted 1:1 ratio (Sensitive: Resistant; Chi-square test, $p>0.05$ )

To test whether the *emb1967* mutations could be suppressed by AtMCRS1 expression, we transformed the heterozygous mutant with a construct for expressing a AtMCRS1-GFP fusion protein under the control of its native promoter. Transformants gave rise to offspring that had AtMCRS1-GFP expressed in the homozygous *emb1967-1* mutation background and produced seeds with normal appearance (Fig. 2). Therefore, we concluded that *AtMCRS1* inactivation was linked and causative to embryo lethality and the AtMCRS1-GFP fusion protein was functional.

### AtMCRS1 is a nuclear protein and does not associate with mitotic microtubule arrays

The functional AtMCRS1-GFP fusion protein allowed us to examine its localization during mitosis, in part because the human MCRS1 localizes to microtubule minus ends in mitotic spindles. We first employed an induced mitotic system in tobacco leaves (Xu et al. 2020). Upon mitotic induction by ectopic Cyclin D expression, we found that AtMCRS1-GFP was detected in nucleoplasm during interphase but did not exhibit a noticeable localization pattern on any specific structures following nuclear envelope breakdown (Supple. Fig. 2).

To capture AtMCRS1 localization in its native cells, we performed live-cell imaging of AtMCRS1-GFP in roots including actively dividing meristematic cells of the rescued plant described above (Fig. 3a). Compared to the wild type root cells that did not have noticeable GFP signals, the rescued plant had the GFP signal confined to nuclei in all root cells at interphase (Fig. 3a). To capture the AtMCRS1-GFP signal with greater details, we also performed immunofluorescence experiments by detecting the protein with anti-GFP antibodies in fixed root tip cells. Prior to nuclear envelope breakdown, again, we found that AtMCRS1 localized to the nucleoplasm of the interphase cell (Fig. 3b). At prophase as highlighted by a mature microtubular preprophase band, the AtMCRS1-GFP signal was detected both in the nuclear region and the cytosol (Fig. 3b). Following the nuclear envelope breakdown, the AtMCRS1-GFP signal was not detected on the mitotic microtubule arrays like the metaphase spindle (Fig. 3b). When the phragmoplast microtubule array was detected after mitosis, AtMCRS1-GFP was not associated with the microtubules either but was detected in the reforming daughter nuclei (Fig. 3b). Therefore, we concluded that AtMCRS1 is a nuclear protein and does not associate with mitotic microtubule arrays.

**Figure 3.**
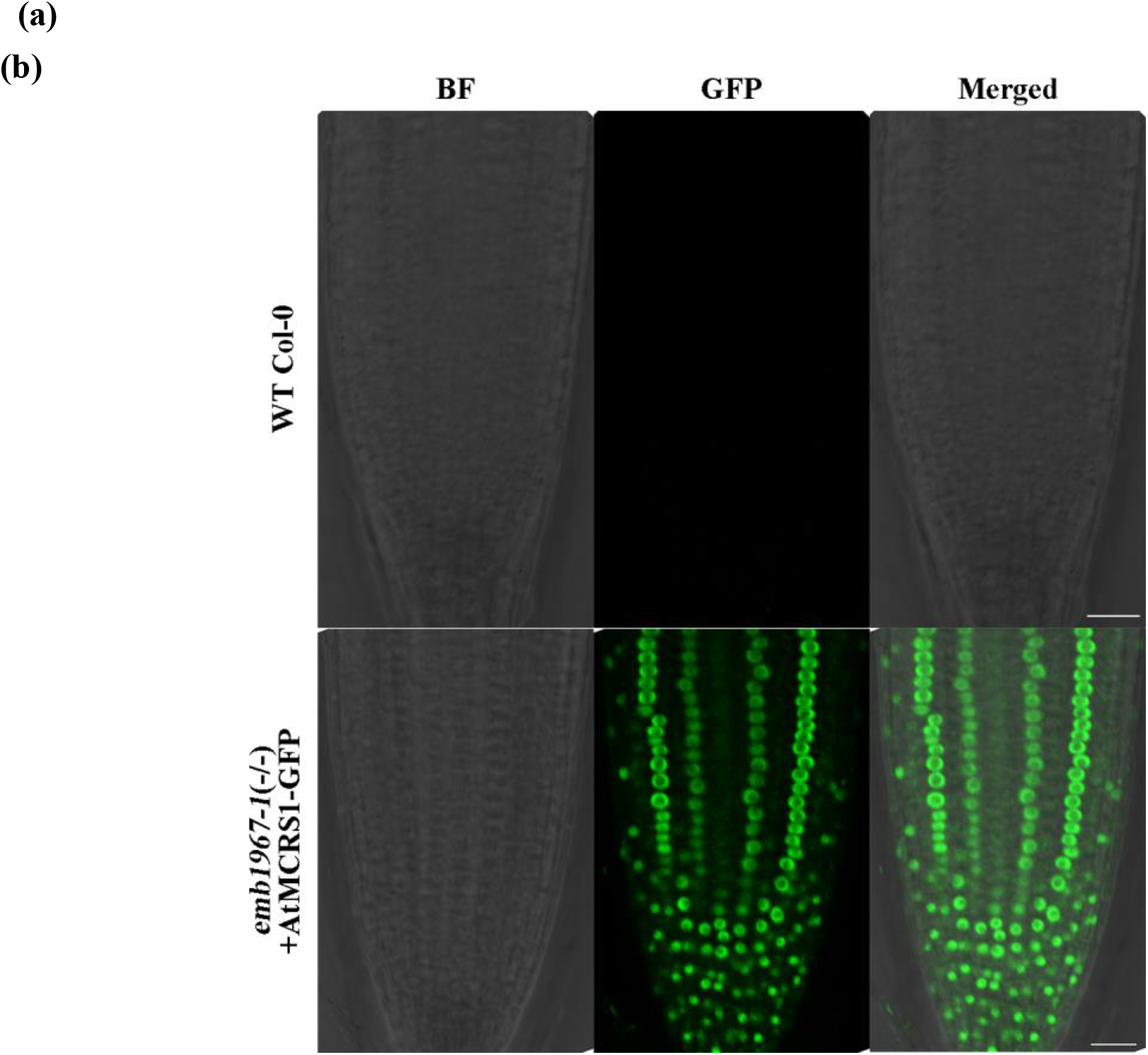
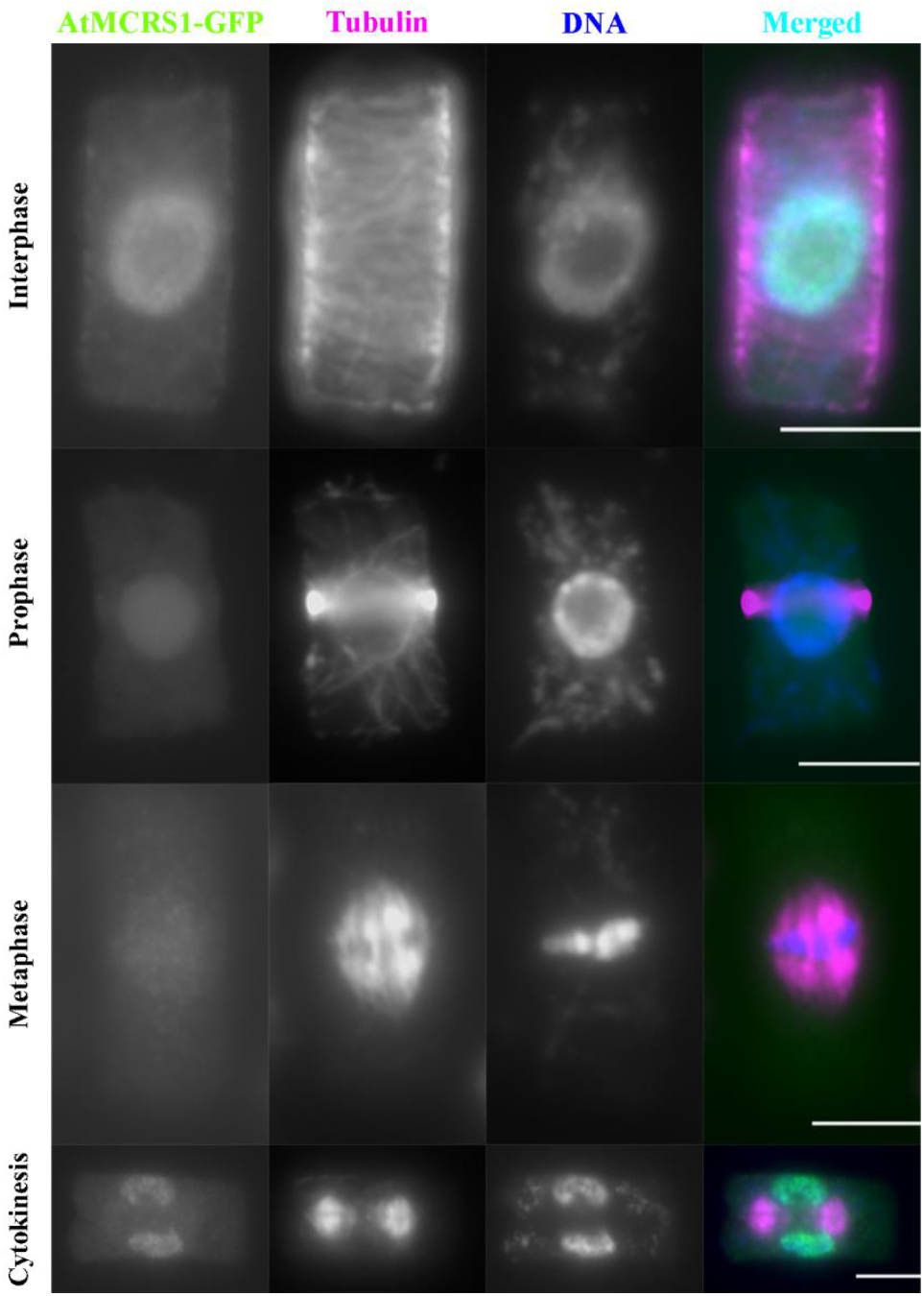
AtMCRS1 is a nuclear protein and does not associate with mitotic microtubule arrays. (a) Live-cell imaging of root cells expressing AtMCRS1-GFP in the homozygous *emb1967-1* background. AtMCRS1-GFP is detected in the nucleoplasm of Arabidopsis root cells. Scale bars= 20 μm; (b) Mitotic cells isolated from roots of plants expressing AtMCRS1-GFP as described above. AtMCRS1-GFP localizes to the nucleoplasm during interphase. At prophase, diffused AtMCRS1-GFP signal is detected in the nuclear region. AtMCRS1-GFP is not noticeably enriched on the metaphase spindle or the phragmoplast. Instead, the GFP signal becomes conspicuous in the reforming nuclei at late cytokinesis. Scale bars=10 μm.

### AtMCRS1 is co-purified with DRMY1

To gain insights into the essential function of AtMCRS1, we aimed to learn what the protein might be associated with in vivo by purifying the AtMCRS1-GFP fusion protein from the rescued plants by anti-GFP affinity purification. When purified proteins were subjected to mass-spectrometry (MS)-assisted peptide identification, AtMCRS1 was detected with 126 total peptides, including 32 unique peptides that covered 47.44% of the polypeptides (Table 3 and Supple. Table 1). We used the nuclear protein BUB3.1 as a reference in the purification to determine proteins specifically detected with AtMCRS1. Mass spectrometry results obtained with GFP BUB3.1-purification (Zhang et al. 2018) did not include AtMCRS1 and vice versa.

**Table 3.** Proteins Identified from AtMCRS1-GFP Pull-down Assay in two replicates.

| Bait | Targets | Unique peptides | Total peptides | Coverage |
| --- | --- | --- | --- | --- |
| AtMCRS1-GFP | MCRS1 | 17/32 | 73/126 | 26.92%/47.44% |
|  | DRMY1 | 23/43 | 67/150 | 33.21%/56.95% |
|  | DP1 | 3/10 | 4/15 | 5.41%/18.36% |
|  | BUB3;1 | 0 | 0 | 0 |
| BUB3;1-GFP | MCRS1 | 0 | 0 | 0 |
|  | DRMY1 | 0 | 0 | 0 |
|  | DP1 | 0 | 0 | 0 |
|  | BUB3;1 | 32 | 443 | 70.00% |

Two independent purification and mass spectrometry experiments resulted in the recovery of a Myb-like protein. This *Development Related Myb-like1* (*DRMY1*) regulates cell expansion and seed production (Wu et al. 2019). DRMY1 was detected with 146 total peptides, including 50 unique ones that covered 56.95% of the DRMY1 polypeptide. Similarly, a paralog of DRMY1, DP1 was also detected with 15 total peptides, including 15 unique peptides that covered 18.36% of the sequence (Table 3). Neither DRMY1 nor DP1 was not detected when BUB3.1 was purified. Our results thus implicated that AtMCRS1 associated with DRMY1 and DP1 *in vivo* in *A. thaliana*.

### AtMCRS1 interacts with DRMY1 directly

Because of the abundant recovery of DRMY1 with AtMCRS1, we predicted that the two proteins likely interact directly. To test this possibility, we applied a technique named CAPPI (Cytoskeleton-based Assay of Protein-Protein Interaction) in living cells (Lv et al. 2017). In this assay, AtMCRS1 was artificially targeted to actin microfilaments after being fused with the 17-amino acid Lifeact peptide (Riedl et al. 2008). The Lifeact-AtMCRS1-GFP fusion protein was targeted to actin filaments when overexpressed in tobacco cells (Fig. 4a). When a DRMY1-TagRFP fusion protein was expressed alone, consistent with the previous reports (Wu et al. 2019), DRMY1 was exclusively detected in the nucleus (Fig. 4a). When Lifeact-AtMCRS1-GFP and DRMY1-TagRFP fusion proteins were co-expressed, however, we found that DRMY1-TagRFP colocalized with Lifeact-AtMCRS1-GFP on actin filaments (Fig. 4b), suggesting that Lifeact-AtMCRS1-GFP targeted DRMY1-TagRFP from the nucleus to the actin filaments. Similarly, Lifeact-AtMCRS1-TagRFP was able to recruit DRMY1-GFP to actin filaments as well. To test the specificity of the recruitment of DRMY1 by AtMCRS1, we co-expressed Lifeact-AtMCRS1-GFP with fluorescent fusion of another nuclear protein, BES1-TagRFP (Lv et al. 2017). BES1-TagRFP remained exclusively in the nucleus while the co-expressed Lifeact-AtMCRS1-GFP appeared both on the actin filaments and in the nucleus (Fig. 4c). To rule out the possibility of the DRMY1 recruitment by the Lifeact peptide, DRMY1-GFP was co-expressed with Lifeact-TagRFP and DRMY1-GFP remained in the nucleus while Lifeact-TagRFP marked actin filaments (Fig. 4d). Therefore, we concluded that AtMCRS1 directly interacted with DRMY1 to result in the redirected localization of the latter in the CAPPI experiment.

**Figure 4.**
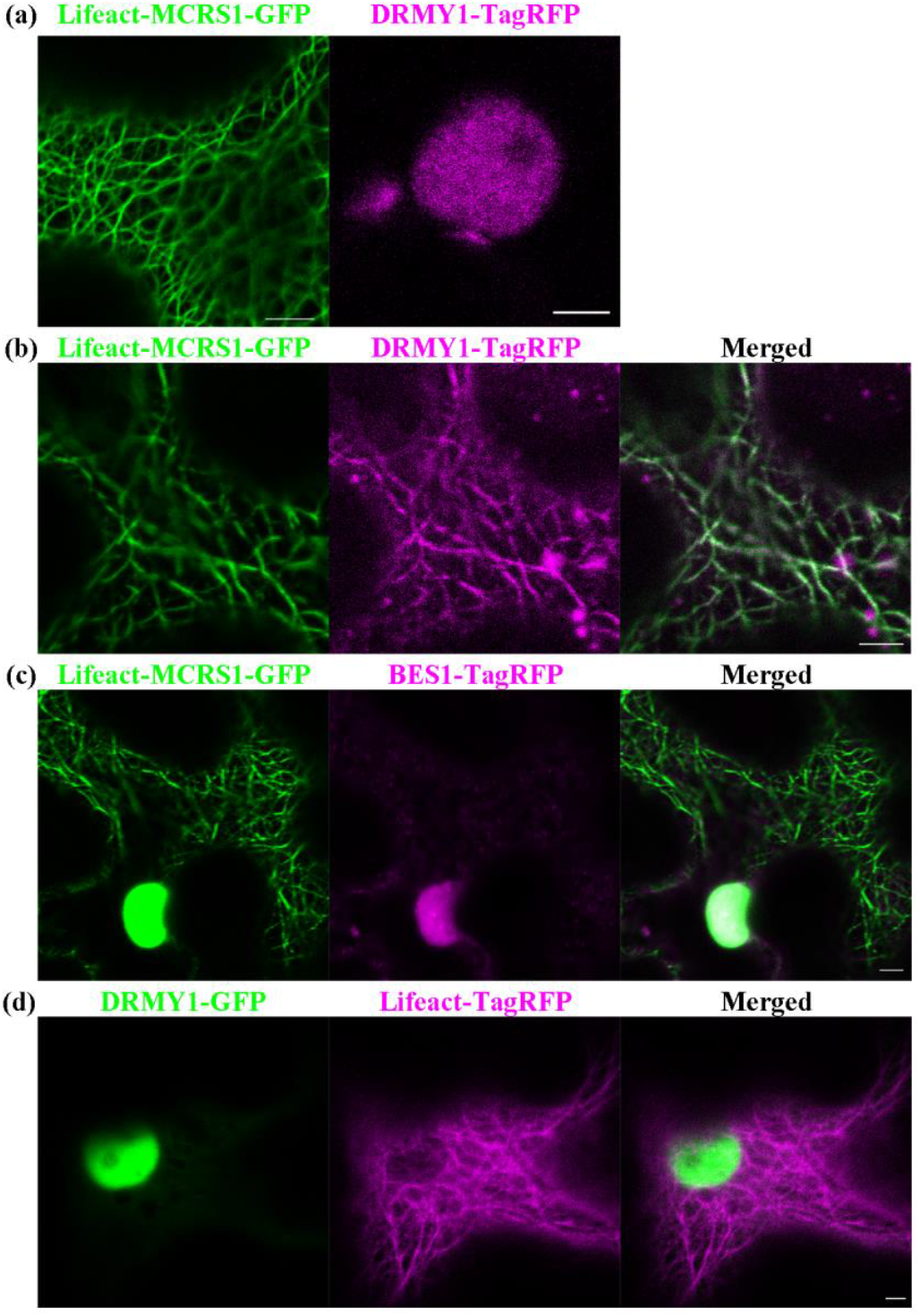
AtMCRS1 interacts with DRMY1. (a) Localizations of Lifeact-AtMCRS1-GFP and DRMY1-TagRFP when they are separately expressed in tobacco epidermal cells. Lifeact-MCRS1-GFP localizes to actin filaments. DRMY1-TagRFP localizes to the nucleus only; (b) Colocalizations of Lifeact-AtMCRS1-GFP and DRMY1-TagRFP fusion proteins when co-expressed in tobacco epidermal cells. Lifeact-AtMCRS1 and DRMY1 colocalizes to the actin filaments, suggesting the interactions between MCRS1 and DRMY1; (c) Localizations of Lifeact-AtMCRS1-GFP and TagRFP fusion of BES1, a non-interacting protein, when co-expressed in tobacco epidermal cells. Lifeact-AtMCRS1-GFP did not change the localization of BES1-TagRFP; (d) Co-expression of Lifeact-TagRFP and DRMY1-GFP in tobacco epidermal cells. Lifeact-TagRFP localizes to the actin filaments, while DRMY1-GFP localizes to the nucleus only. Scale bars=5 μm.

## Discussion

In this study, we characterized the function of the evolutionarily conserved AtMCRS1 in *A. thaliana* and reported that AtMCRS1 is an essential nuclear protein that interacted with the developmentally important DRMY1 protein so that is hypothesized to function as a transcription regulator. Unlike its human counterpart, however, AtMCRS1 did not exhibit a cell cycle-dependent, microtubule-associated localization pattern so that did not play a role in mitosis as indicated in the mutant as well.

### AtMCRS1 functions as a potential transcription regulator

Our results demonstrated the direct interaction between AtMCRS1 and DRMY1, which introduced a previously unknown mode of interaction for this evolutionarily conserved protein. Among proteins that are associated with the human MCRS1 protein, many reside in the nucleolus and are implied in gene expression associated with ribosome biosynthesis or sequestration of transcription factors to the nucleolus (Lin and Shih 2002; Shimono et al. 2005). In general, MCRS1 is found to transcriptionally promote cell proliferation and stress responses in animals by directly interacting with a wide collection of proteins through different functional domains (Huang et al., 2022). Both our protein purification and CAPPI assay revealed that DRMY1 acts as a bona fide partner of AtMCRS1 *in vivo* in *A. thaliana*, suggesting that MCRS1 functioned as a major regulator of DRMY1 function. It has been reported that DRMY1 is involved in gene expression programs associated with GO terms like “stress response”, “cell wall modification”, and “hormone response” and its loss leads to severe reduction of seed production attributed to severe defects in cell expansion (Wu et al. 2019; Zhu et al. 2020). In fact, it was found that DRMY1 functions as a positive regulator of cell wall biosynthesis/remodeling- and ribosome biosynthesis-related gene expression and represses of genes related to ABA and ethylene signaling (Wu et. Al., 2019). It would be interesting to test whether AtMCRS1 was also involved in these processes. Unfortunately, such a test would rely on the generation of hypomorphic mutants because of the essential function of the protein in *A. thaliana*. Alternatively, DRMY1 might function as a regulator of AtMCRS1 in the nucleus. Unlike the animal MCRS1 proteins, AtMCRS1 does not possess an obvious nuclear localization signal. Therefore, it is possible that it may be dependent on its interacting partner(s) to achieve the nuclear localization. Nevertheless, it will be interesting to test whether these two proteins act together in transcription regulation of downstream genes.

### AtMCRS1 does not a function in mitosis

Because of the reported MCRS1 function in stabilizing the minus ends of acentrosomal microtubules in mitotic spindles (Meunier and Vernos 2016), such a role would be highly applicable for plant mitosis simply because flowering plant do not produce the centrosome structure. Therefore, it was surprising for us to learn that the AtMCRS1 protein was totally dispensable for mitotic cell division during gametophyte development. In fact, the lack of association with mitotic microtubule arrays was consistent with no impact of AtMCRS1 inactivation on mitotic cell divisions associated with gametogenesis. We reasoned that the lack of such a function known to human MCRS1 is probably because of the absence of the MCRS1 interacting partners that are necessary for MCRS1 to decorate microtubule minus ends. In human cells, MCRS1 association with microtubules is brought about by the interaction with the KANSL protein complex which includes the KANSL3 as a microtubule minus end-associated protein (Meunier et al., 2015). When the human KANSL3 polypeptide sequence was compared to sequences of the proteins in *A. thaliana*, no obvious homologous protein was identified. The absence of a plant homolog of KANSL3 perhaps attributed to the lack of AtMCRS1 association with spindle microtubules.

## Materials and Methods

### Multiple sequences alignment

The amino acid sequences of HsMCRS1 and its closest land plant homologs were acquired from NCBI (https://www.ncbi.nlm.nih.gov/), TAIR (https://www.arabidopsis.org/), and Phytozome (https://phytozome-next.jgi.doe.gov/). The accession numbers of the proteins included are: HsMCRS1 (Q96EZ8.1), AtMCRS1 (NP_001325804.1), OsMCRS1 (XP_015632662.1), PpMCRS1(XP_024389733.1), SmMCRS1 (XP_024518220.1), and MpMCRS1 (PTQ39771.1). The sequences were aligned using Clustal Omega (Madeira et al. 2019) with default settings and the results were exported using Jalview. The illustrations of MCRS1 domain architecture were generated using IBS 1.0 (Liu et al. 2015).

### Plant Materials and Growth Conditions

The Arabidopsis plants were grown in a growth chamber at 21□ under a 16-hr light/8-hr dark cycle with 70% relative humidity. The tobacco plants (*Nicotiana benthamiana*) were grown in a growth chamber at 24□ under a 10-hr light/14-hr dark cycle.

For live-cell imaging and immunolocalization experiments, the seedlings were germinated on solid medium with half-strength MS (Murashige & Skoog) salt mixture and 0.8% phytagel.

Both heterozygous *emb1967-1* and *emb1967-2* mutant seeds were acquired from the Arabidopsis Biological Research Center at Ohio State University with the stock numbers of CS16040 and CS24096. The *emb1967-1* mutation was detected by the primer pair of 16040RP (5’-CGT CAT CAC CCA GCC GTA TGA TCG C-3’) and GLB3 (5’-TAG CAT CTG AAT TTC ATA ACC AAT CTC GAT ACA C-3’) while the wild type locus was detected by 16040RP and 16040LP (5’-ATA CTT GAC ATG GAC TTG GAA CCT G-3’). Similarly, the *emb1967-2* mutation was detected by the primer pair of 24096RP (5’-CAG ACA AGC TGA AAG AAG AAA CAC-3’) and GLB3, and the wild type allele by 24096LP (5’-GGG GCT CTT GCT CAA GTT GTT CCG-3’) and 24096RP. To verify the recovery of homozygous *emb1967-1* mutant expressing AtMCRS1-GFP, the primers 16040RP and MCRS1_veri (5’-AAC ACG CAC TCT TGA TCT GGT CTC-3’) were used as they only rendered a PCR product from the wild type gene but not the fusion construct.

### Plasmid construction

An *AtMCRS1* fragment was amplified by the Phusion DNA polymerase (ThermoFisher) from the *A. thaliana* genomic DNA using the primer pair III54350Fb1 (5’-GAC AAG TTT GTA CAA AAA AGC AGG CTC CGC ACG AAT CAA CAA AAT TAG GTC AC-3’) and III54350Rb2 (5’-GAC CAC TTT GTA CAA GAA AGC TGG GTC GTT CAC TTT CCC TCT TCT CTT CAG G-3’). The resulting product was inserted into pDONR211 by using BP clonase (ThermoFisher) to generate pENTR-AtMCRS1. pENTR-AtMCRS1 was then assembled in pGWB4 (Nakagawa et al. 2007) by using LR clonase (ThermoFisher) to generate the expression construct pGWB4-AtMCRS1 for producing the AtMCRS1-GFP fusion protein.

To prepare expression constructs for the CAPPI assay, cDNA fragments of AtMCRS1 and DRMY1 were amplified from an *A. thaliana* cDNA library by Phusion using primer pairs of GibMCRS1F (5’-GTT TGA GTC TAT CTC CAA AGA GGA GGG AGC TGG AGC TAT GGG GGC TCT TGC TCA AG-3’) and GibMCRS1R (5’-GTC GGC GCG CCC ACC CTT GTT CAC TTT CCC TCT TCT CTT CAG GT-3’); and GibDRMY1F (5’-ATG GGA ACC AAT TCA GTC GAC TGG ATG GCG TAT TAT GCT GAA GCA AAG A-3’) and GibDRMY1R (5’-CTT TGT ACA AGA AAG CTG GGT CTA GAT ACA ACT CCT TCA GTC CGG TCC-3’), respectively. The MCRS1 fragment was inserted into a pENTR-Lifeact plasmid (Lv et al. 2017) at the EcoRI-BamHI sites and the DRMY1 fragment into pENTR4 at EcoRI-BamHI sites by Gibson Assembly to produce pENTR-Lifeact-AtMCRS1 and pENTR-DRMY1, respectively. pENTR-Lifeact-AtMCRS1 and pENTR-DRMY1 were then assembled into pGWB5 and pGWB660 (Nakamura et al. 2010)by LR clonase to generate pGWB5-Lifeact-AtMCRS1 (Lifeact-AtMCRS1-GFP), pGWB5-DRMY1 (DRMY1-GFP), pGWB660-Lifeact-AtMCRS1 (Lifeact-AtMCRS1-TagRFP), and pGWB660-DRMY1 (DRMY1-TagRFP). BES1-TagRFP and Lifeact-TagRFP constructs were previously constructed (Lv et al. 2017).

### Stable transformation in *Arabidopsis thaliana*

The *emb1967-1* heterozygous mutant plants were transformed with the pGWB4-MCRS1 plasmid by the floral dip method using *Agrobacterium tumefaciens* strain GV3101. GV3101 cells carrying the plasmid were grown in liquid LB medium supplemented with corresponding antibiotics at 28□ for 2 days. Bacterial cells were pelleted by centrifugation at 5000 rpm for 10 min under 4□ and resuspended in 5% sucrose solution containing 0.05% Silwet L-77 for dipping. The transformants were selected on solid medium with half-strength MS (Murashige & Skoog) salt mixture and 0.35% phytagel supplemented with 20 μg/ mL hygromycin.

### Transient expression in *Nicotiana benthamiana*

Transient expression experiments were performed in leaves of *N. benthamiana* by agrobacterial infiltration. GV3101 bacteria carrying the expression vectors were grown in LB medium supplemented with corresponding antibiotics at 28□ for 2 days. Bacterial cells were pelleted by centrifugation at 3000 rpm for 10 min under room temperature and resuspended in infiltration buffer (10 mM MES pH 5.7, 10 mM MgCl_2_, 150 μM acetosyringone). Cultures carrying different vectors were mixed and adjusted to a final OD_600 nm_ of 1.0 for each culture for co-infiltration. These cultures were then mixed with *A. tumefaciens* C58C1 (pCH32-35S::p19, adjusted to a final OD_600 nm_ of 0.7) and incubated for 3 hr at room temperature. The combined bacterial suspensions were infiltrated into leaves of 4-week-old *N. benthamiana*. The leaf samples were observed under microscope 2 days post infiltration.

### Immunolocalization and fluorescence microscopy

Root tips of Arabidopsis were fixed for 45 min in 4% formaldehyde in PME (50mM Pipes buffer, pH 6.9, 1mM MgSO_4_, and 5 mM EGTA). After partially digested for 23 min in 1% cellulase and 0.1% pectolyase in PME, root tip cells were released by gentle squashing onto coated glass slides. Following sequential treatment with 0.5% Triton X-100 for 15 min and methanol at -20□ for 10 min, the cells were processed for immunofluorescence staining. Stained cells were observed under an Eclipse 600 microscope equipped with 60X Plan-Apo and 100X Plan-Fluor objectives (Nikon) and images were captured by an OptiMOS sCMOS camera (Photometrics) controlled by the µManager software package (https://micro-manager.org/) (Edelstein et al. 2014).

In live-cell imaging experiments, root tips or tobacco leaf segments were mounted in water before being placed an Axio Observer inverted microscope equipped with the LSM710 laser scanning confocal module (Carl Zeiss). Cells were observed using 40X C-Plan (water) objectives, and the GFP fusions were excited by 488 nm argon laser and TagRFP by 561 nm argon laser. Emitted fluorescence signals were collected according to manufacturer’s instruction. Images were acquired by the ZEN software package and processed in Image J (https://imagej.nih.gov/ij/).

### Protein affinity purification and peptide identification by mass-spectrometry

Proteins were extracted from the transgenicAtMCRS1-GFP plants. Briefly, flower buds were collected and frozen in liquid nitrogen prior to being ground using a mortar and a pestle. Extraction buffer (50 mM Tris-HCl, pH 8.0, 150 mM NaCl, and 1% Triton X-100) was added to the powder and incubated at 4□ for 1h. The supernatant was collected after centrifugation at 15,000*g* and filtration through a 0.45 μm Millex HV filter (Millipore). AtMCRS1-GFP was purified with anti-GFP antibody-conjugated magnetic beads (Miltenyi Biotec) according to the manufacturer’s instruction. Bound proteins were eluted by the SDS-PAGE sample buffer and subjected to separation in 7.5% polyacrylamide gels. Gels were stained with silver nitrate or Coomassie Blue.

Peptide identification by mass spectrometry was carried out at the Taplin Mass Spectrometry Facility of Harvard Medical School. Following mass-spectrometric analysis of peptides resulted from trypsin digestion, they were identified by referencing to the *A. thaliana* proteome annotated by TAIR (https://www.arabidopsis.org/).

### Quantification and statistical analysis

Microsoft Excel was used to perform Chi-square test in Table 1 and Table 2.

## Data Availability

The data underlying this article are available in TAIR (The Arabidopsis Information Resource): https://www.arabidopsis.org/.

## Funding

This work was supported by the NSF grant MCB-1920358 and the U. S. Department of Agriculture (USDA)-the National Institute of Food and Agriculture (NIFA) under an Agricultural Experiment Station (AES) hatch project (CA-D-PLB-2536-H).

## Acknowledgments

We thank Dr. Tsuyoshi Nakagawa at Shimane University in Japan for generously sharing the pGWB plasmids.

## Author Contributions

BL conceptualized the project. All authors were involved in experimental designs. HH and ML performed the experiments and collected the data. All authors participated in data analysis. HH wrote and BL revised the manuscript.

## Disclosures

The authors have no competing interests.

## Notes

### Competing Interest Statement

The authors have declared no competing interest.

