## Supplemental figures for "The Microspherule protein 1(MCRS1) homolog interacts with the Myb-like transcription factor DRMY1 and is essential for embryogenesis in *Arabidopsis thaliana*"

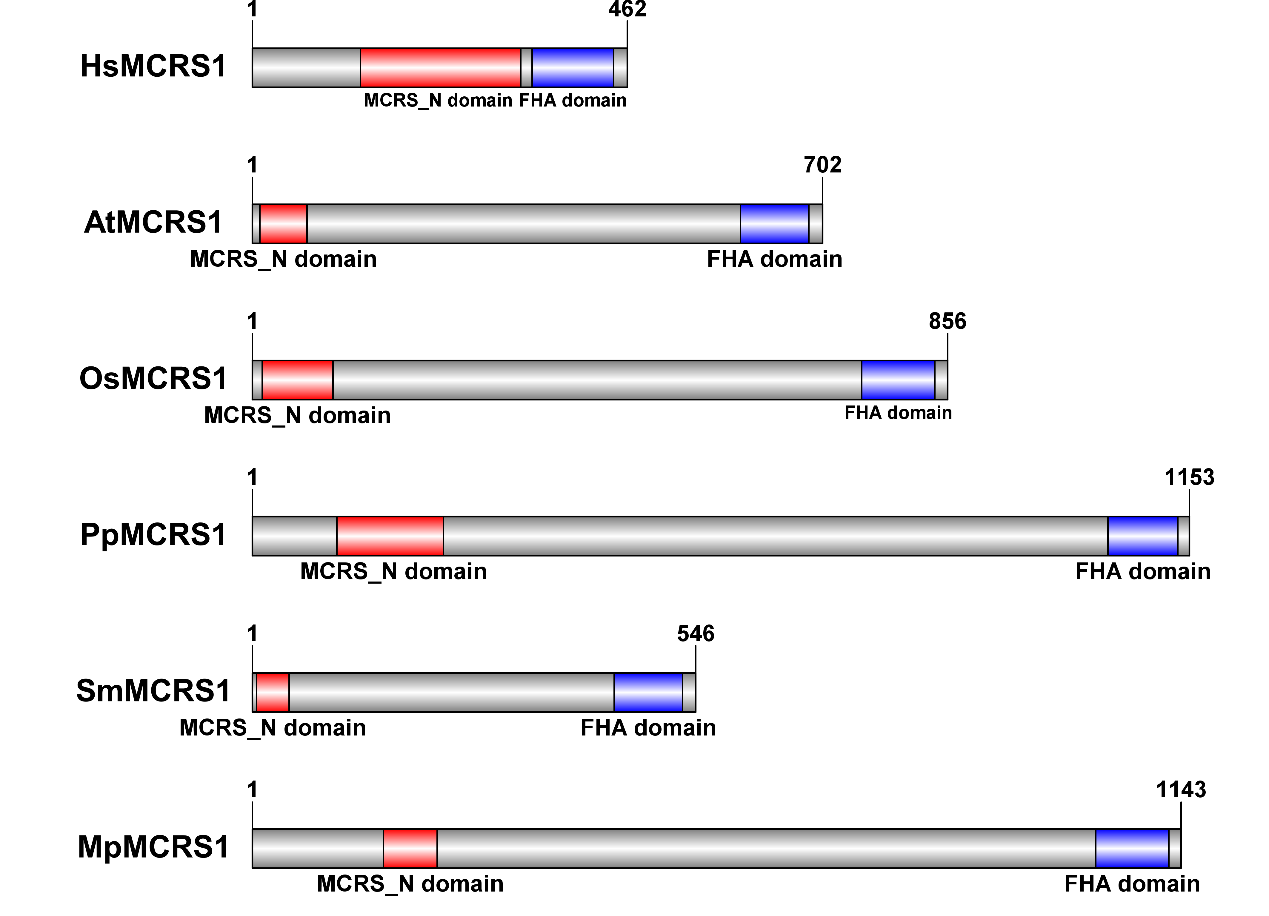
**(a)**

**(b)**

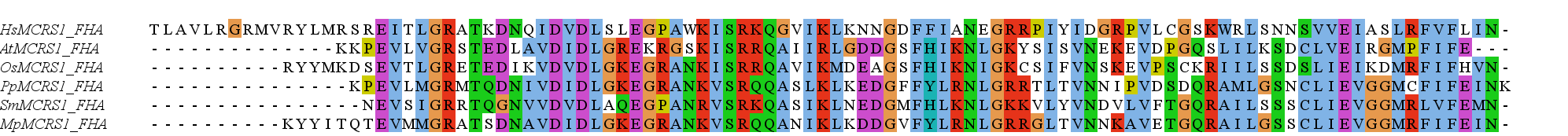

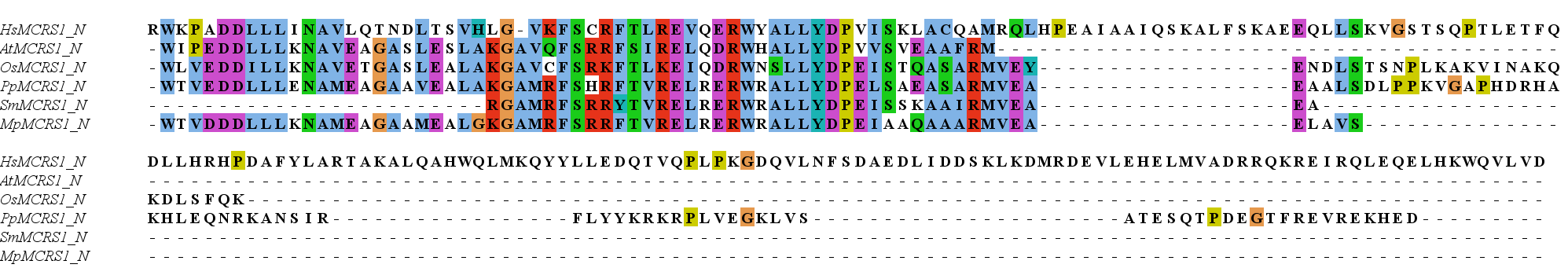

**Supplementary Figure 1 Homology between HsMCRS1 and homologs in land plants.** (a) Domain architectures of HsMCRS1 and the plant homologs. The predicted MCRS_N domains are labeled in red and FHA domains in blue; (b) Sequence alignments of the MCRS_N domains and FHA domains. MCRS_N and FHA domain sequences of HsMCRS1 and the plant homologs were obtained from NCBI and aligned with Clustal X. The conserved amino acids are labeled in colours. Hs: *Homo sapiens*; At: *Arabidopsis thaliana*; Os: *Oryza sativa*; Pp: *Physcomitriella patens*; Sm: *Selaginella moellendorffii*; Mp: *Marchantia polymorpha*.

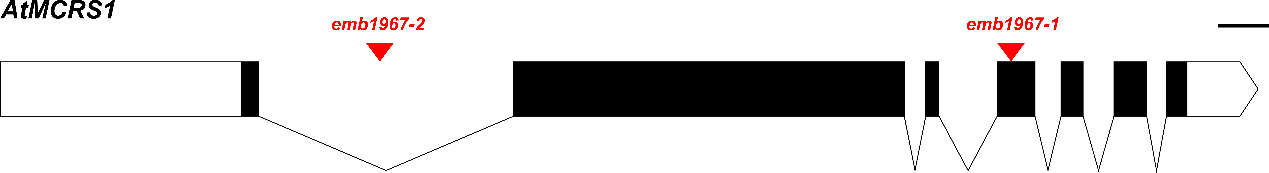

**Supplementary Figure 2 Gene model of *AtMCRS1* (AT3G54350).** Two T-DNA insertion mutations were identified. *emb1967-1* (CS16040) has an insertion in Exon 4. *Emb1967-2* (CS24096) has an insertion in Intron 1.

**Supplementary Table 1 Peptides Identified from AtMCRS1-GFP Pull-down Assay.** The recovered peptides are highlighted in yellow.

**DRMY1 (AT1G58220.1)**

```
  1 MVDNSNNKKR KEFISEADIA TLLQRYDTVT ILKLLQEMAY YAEAKMNWNE
 51 LVKKTSTGIT SAREYQLLWR HLAYRDSLVP VGNNARVLDD DSDMECELEA
101 SPGVSVDVVT EAVAHVKVMA ASYVPSESDI PEDSTVEAPL TINIPYSLHR
151 GPQEPSDSYW SSRGMNITFP VFLPKAAEGH NGNGLASSLA PRKRRKKWSA
201 EEDEELIAAV KRHGEGSWAL ISKEEFEGER TASQLSQRWG AIRRRTDTSN
251 TSTQTGLQRT EAQMAANRAL SLAVGNRLPS KKLAVGMTPM LSSGTIKGAQ
301 ANGASSGSTL QGQQQPQPQI QALSRATTSV PVAKSRVPVK KTTGNSTSRA
351 DLMVTANSVA AAACMSGLAT AVTVPKIEPG KNAVSALVPK TEPVKTASTV
401 SMPRPSGISS ALNTEPVKTA VAASLPRSSG IISAPKVEPV KTAASAASLP
451 RPSGMISAPK VEPVKTTASV ASLPRPSGII SAPKAEPVKT AASAASSPRP
501 SGMISAPKVE SVKTTASMPR PSGIISAPKA ELVKSAASAA SLPCTSGIIS
551 SPKAELVKSA ASAASFPRPS SMLSAPKADP VKIVPAAATN TKSVGPLNLR
601 HAVNGSPNHT IPSSPFTKPL HMAPLSKGST IQSNSVPPSF ASSRLVPTQR
651 APAATVVTPQ KPSVVAAATV VTPQKPSVGA AATVVTPQKP SVGAAANVVT
701 PQKPSVGSAA TVVTPQKPSV GAAVTVTSKP VGVQKEQTQG NRASPLVTAT
751 LPPNKTIPAN SVIGTAKAVA AKVETPPSLM PKKNEVVGSC TDKSSLDKPP
801 EKESTTTVSP LAVAATKSKP KDEATVTGTG LKEL
```

**DP1 (AT1G58220.1)**

1 MVANNNTSSN RRKRIITEGD IATLLLRYDM ETILRMLQEI SYCSETKMDW
 51 NALVKKTTTG ITNAREYQLL WRHLSYRHPL LPVEDDALPL DDDSDMECEL
101 EASPAVSHEA SVEAIAHVKV MAASYVLSES DILDDSTVEA PLTINIPYAL
151 PEGSQEPSES PWSSRGMNIN FPVCLQKVTS TEGMNGNGSA GISMAFRRKR
201 KRWSAEEDEE LFAAVKRCGE GNWAHIVKGD FRGERTASQL SQRWALIRKR
251 CHTSTSVSQC GLQGTEAKLA VNHALSLALG NRPPSNKLAI GTSSRRSFPA
301 NSSIYVITED ALVWLPLACL NQKLAYLFNC GLMPTTSSCT ITETEANGGS
351 SSQGQQQSKP IVQALPRAGT SLPAAKSRVV KKTTASSTSR SDLMVTANSV
401 AAAACMGDVL TAASGRKVEP GKTDAPRVPK TKPVKHASTV CMPQPSGSLS
451 MPKVEPGTSV AASIRSLANG KLKPVMASSS SNKPPLIAPR SEGSSMLSAS
501 APLASLSRIV SNQRVFAGSV PATEIVTCKP DGGQKGQARG NEASSSAAIQ
551 PHQITSRNLE ISQGKQATQA QSPNLLPRKV PVVRTAVHCA TNQKLMDKPS
601 DQTVVPIRGA GSQSKAKGEV NSKVGPVIKV SSVCGKPLEV ATVAGTGQGV

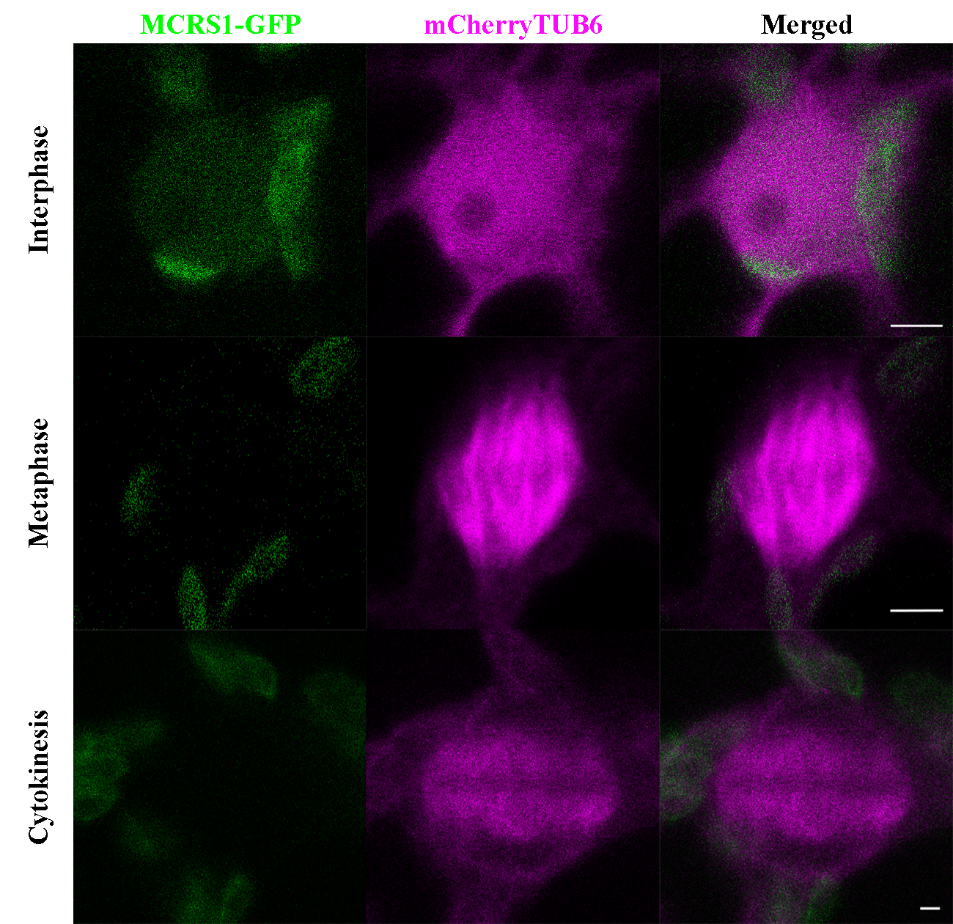

**Supplementary Figure 3 Expression of AtMCRS1-GFP in tobacco leaf epidermal cells induced to enter mitosis.** The GFP signal is detected in the nucleus in interphase. The GFP signal is not enriched on the mitotic spindle or the phragmoplast. Scale bars=5 μm.
